# Database of recurrent mutations (DORM), a web tool to browse recurrent mutations in cancers

**DOI:** 10.1101/2022.11.21.517363

**Authors:** Deepankar Chakroborty, Ilkka Paatero, Kari J. Kurppa, Klaus Elenius

## Abstract

Advances in sequencing technologies have facilitated the genetic characterization of large numbers of clinical cancer samples, leading to accumulation of extensive amounts of data. While potentially very useful for directing research and for clinical decision making, the increasing quantity of data generates challenges in its optimal management, and translation to informing clinical and research questions. Here, we present <u>D</u>atabase <u>O</u>f <u>R</u>ecurrent <u>M</u>utations (DORM), a database listing recurrent mutations (tissue-agnostic population frequency > 1) identified from cancer samples analyzed with whole genome or whole exome sequencing. The DORM database is a fast and feature-rich database supporting searching for several proteins, amino acid substitutions as well as queries using regular expressions.

## INTRODUCTION

The fast-paced development of next-generation sequencing (NGS) technology and its use to study cancer specimens has led to an accumulation of large amounts of data and establishment of expansive databases that have propelled the discovery of predictive and therapeutic biomarkers for various cancers (Campbell *et al*. 2020). Large-scale sequencing efforts have pinned somatic mutations as the most common cause of human cancers (Martincorena and Campbell 2015). Mutations in several oncogenes are well-characterized driver events in various cancers, e.g. mutations in KRAS G12 residue in pancreatic and lung cancer (Hong *et al*. 2020), BRAF V600 in melanoma (Hauschild *et al*. 2012), and the EGFR L858 in lung cancer (Lynch *et al*. 2004; Paez *et al*. 2004). Despite their frequent observations in the clinic, these hotspot mutations actually make up a small proportion of all the cancer-associated mutations and there are a large number of recurrent, but “non-hotspot” mutations (Chang *et al*. 2016).

Databases presenting cancer-associated mutations like COSMIC (https://cancer.sanger.ac.uk) (Tate *et al*. 2019), the ICGC data portal (https://dcc.icgc.org) (Zhang *et al*. 2011), AACR GENIE (https://genie.cbioportal.org) (The AACR Project GENIE Consortium 2017), and cBioportal (https://www.cbioportal.org/) (Cerami *et al*. 2012; Gao *et al*. 2013) present a well-designed interface that provides access to rich data. However, by design, these databases with comprehensive information use a significant amount of bandwidth as well as require multiple steps to access to key pieces of information, like the frequency of mutations and the affected amino acid residues. Especially, calculation of the latter requiring manual processing of the data, as there is no direct way to retrieve this information from any of these large databases.

We sought to solve these shortcomings and built a database of recurrent mutations using the large COSMIC cancer registry as a model. Our goal with this project was to develop and deploy a fast and lightweight web-tool to give the user a quick-and-easy way to check the status of a particular mutation of interest in cancer samples in an easy-to-understand format. In addition to direct time-savings, we believe initiatives like ours, help further cancer research and its global outreach by improving accessibility to well-summarized information. Moreover, we hope that our open-source framework enables applications to other public cancer registries and diversification to other frontiers of healthcare.

## MATERIALS AND METHODS

### Website and web server

The DORM database is accessible at https://eleniuslabtools.utu.fi/tools/DORM/Mutations/, and all requests to the server are handled by an NGINX reverse-proxy (https://nginx.org/) that encrypts the traffic between our server and the end-user’s web-browser. The connection is encrypted using the latest Transport Layer Security (TLS) cryptographic protocol 1.3 (Rescorla 2018) and an industry standard 256-bit Advanced Encryption Standard (AES-256) (National Institute of Standards and Technology 2001). Additionally, as a fallback, the server of DORM also supports connections over TLS 1.2 to support legacy hardware and browsers. The landing page website and the documentation is built using HTML5, CSS and JavaScript. The web tools are built using Shiny (Chang *et al*. 2021) and R (R Core Team 2018). These services are hosted on a virtual private server at the premises of University of Turku, Turku, Finland. The source code for deploying DORM as an R Shiny app is available at https://github.com/dchakro/DORM_Mutations repository.

### Hardware

#### Database processing & analysis

Apple iMac (early 2013) equipped with Intel Core i5 CPU (4 cores – 3.2 GHz), 24 GB DDR3 RAM, 500 GB SSD running macOS Catalina 10.15.

#### Server

Virtual private server (KVM virtualization) with Intel(R) Xeon(R) Gold 5120 CPU (1 core – 2.20 GHz), 6 GB ECC RAM, 100 GB HDD running Ubuntu 18.04 LTS.

#### Web performance testing

Apple MacBook Pro (early 2015) equipped with an Intel Core i5 CPU (2 cores – 2.7 GHz), 8 GB DDR3 RAM, 500 GB SSD running macOS Catalina 10.15. The device was connected via a 5 GHz Wi-Fi router to the public ISP (i.e., outside the network where the DORM database is hosted) over a 100 Mbps fiber optic broadband connection.

### Data and processing of data

Data were acquired from COSMIC release v95 (released November 24, 2021 https://cancer.sanger.ac.uk) as a GNU zip (GZIP) archive of the tab-delimited text file with all mutations identified from genome-wide screens (includes data from whole genome sequencing, and whole exome sequencing). The samples from targeted screens were excluded to ensure our analysis is free from selection bias and so that, for a particular tissue, the frequency of mutations between different proteins can be compared directly.

#### Pre-processing

The decompressed data (16.03 GB) is processed using the “awk” programming language (Aho, Kernighan and Weinberger 1988) to select relevant columns (named, Gene name, Sample name, Primary site, Primary histology, Genome-wide screen, Mutation CDS, Mutation AA). This step reduces the size of the data matrix by ≈80%, thereby, decreasing the computation time and computational resource requirements for the downstream analyses. The selected columns were read in R by using the `data.table::fread()` function (Dowle and Srinivasan 2021). The complete database was stored as standard R object in the .RDS file format, with a notable difference: instead of `saveRDS` from R “base”, which uses serialized compression, parallelized GZIP (pigz: https://zlib.net/pigz/) was used for compression – decompression. This enabled usage of multiple CPU threads to speed up the read-write operations, and in our case was limited by disk I/O. The functions for reading-writing R objects in .RDS files using parallelized compression-decompression are described in this R script.

#### Filtering

The duplicate entries for mutations attributed to ENSEMBL transcripts (n = 30.8 × 10^6^) were removed (Supplementary Figure S1). Mutations with unknown consequences on the protein level (n = 8.8 × 10^6^) were removed, leaving 6.4 × 10^6^ coding alterations. From these, silent mutations (n = 1.5 × 10^6^) i.e., nucleotide substitutions leading to no changes at the amino acid level (this phenomenon happens due to codon degeneracy (Watson *et al*. 2007)) were removed (Supplementary Figure S1). To retain only unique entries, a mutation ID was created using the sample name, protein name and the amino acid change. Samples with duplicate mutation IDs (n = 78,569) were removed (Supplementary Figure S1). The filtered database with unique coding mutations (n = 4.8 × 10^6^) were stored as a parallelized GZIP .RDS file, enabling faster load times. Searching and parsing of the text was done using the ‘stringi’ R package (Gagolewski 2021).

#### Processing

Mutations with single occurrences (i.e., frequency = 1) were removed (n = 2.9 × 10^6^) from the list of unique coding mutations, as they are not part of the pool of recurrent mutations. For each mutation, its cumulative frequency of occurrence, as well as its frequency in cancers of various tissues, was calculated and compiled into a table. The table was sorted by mutation frequency (total number of samples across all cancers) and then stored as a parallelized-GZIP .RDS file.

#### Updates

Since 2004, marking the release of COSMIC v1, the dataset has been updated on average four times per year (range: 11 releases in 2006 and two releases in 2020). The COSMIC data releases need to be acquired from (https://cancer.sanger.ac.uk), then our optimized pipeline can be run with a shell script that automates the processing and generation of the underlying database for DORM.

### Benchmarking and testing performance

To evaluate the performance of different code blocks, the ‘microbenchmark’ R package (Mersmann 2021) was used to gather data. The data were graphically represented using Graphpad Prism 9. Statistical testing comparing multiple groups was performed using Brown Forsythe and Welch ANOVA test and correction for multiple testing was done by controlling the false discovery rate using the two-stage step-up method of Benjamini, Krieger and Yekutieli (Benjamini, Krieger and Yekutieli 2006) in Grahpad Prism 9. Statistical testing comparing two groups of observations was done using Welch’s t-test in Grahpad Prism 9. The code blocks used for testing and benchmarking their performance is available at https://github.com/KE-group/DORM-2022 repository.

The performance of the websites hosting the databases was measured on Google Chrome (v. 97.0.4692.99) with Google Lighthouse (v. 8.5.0) (available in Chrome DevTools). Lighthouse (https://github.com/GoogleChrome/lighthouse) is an open-source tool for automated auditing and assessing performance metrics. A search for EGFR mutations was done on the five databases (DORM, COSMIC, ICGC, cBioPortal and AACR GENIE), and links (Supplementary Table 1) to those individual searches were used to test the performance of the databases. This was done to discount the varying duration required to do the same search on the four databases. Lighthouse 8 produces a performance score which is a weighted average of First Contentful Paint (10%, marks the time at which the first text or image is painted), Speed Index (10%, shows how quickly the contents of a page are visibly populated), Largest Contentful Paint (25%, marks the time at which the largest text or image is painted), Time to interactive (10%, the amount of time it takes for the page to become fully interactive), Total Blocking Time (30%, measures the total amount of time that a page is blocked from responding to user input), and Cumulative Layout Shift (15%, measures the unexpected movement of page content). The JSON data in the lighthouse reports was parsed using the ‘jsonlite’ R package (Ooms 2014) and tabulated in R. The data were graphically represented using Graphpad Prism Statistical testing comparing multiple groups was performed either using Brown Forsythe and Welch ANOVA test or Kruskal-Wallis test. Correction for multiple testing was done by controlling the false discovery rate using the two-stage step-up method of Benjamini, Krieger and Yekutieli (Benjamini, Krieger and Yekutieli 2006) in Grahpad Prism 9.

## RESULTS

### Optimizing R code for speed and efficiency

In order to maximize the efficiency throughout our pipeline, we benchmarked common workflows that can be used to resolve the computation bottlenecks. There are several packages for reading data into R, and we discovered that `data.table::fread()` function offered the best performance in reading both small and big (tested with 10^5^ rows) tables. In our tests, `data.table::fread()` was faster (*q* < 0.0001) than `base::read.table()` in reading files containing 10^3^ and 10^5^ rows (Supplementary Figure S2 A-B). As intended by the developers of the ‘readr’ and ‘vroom’ package, their individual functions for reading TSV files were faster than fread for large files (*q* < 0.001), but, they were slower by a factor of 16-20 for smaller files (*q* < 0.0001) (Supplementary Figure S2 A-B). Furthermore, the data.table implementation of generating a frequency table was more efficient (*q* < 0.0001) than alternatives like `base::table()` and `plyr::count()` (Supplementary Figure S2 C). Consequently, we used `data.table` as the background framework for reading and managing tabular data throughout our analysis pipeline, as well as in the backend of the R Shiny web tool for DORM.

Filtering a large database requires extensive use of search, and the search-replace functionality is required for parsing and cleaning up data fields. We found that alternatives from ‘stringi’ were faster than their counterparts in R base (*P* < 0.0001) (Supplementary Figure S2 D) in carrying out these operations. Furthermore, in comparison to `base::grep()`, we observed considerable improvements in speed (500-fold reduction, *q* < 0.0001) by using GNU-grep for searching the data (stored on disk) server-side when a query was received by the R Shiny web tool (Supplementary Figure S2 E). Therefore, the fastest approach to process the search queries (entered by a user in the interface), was to execute the search using GNU-grep on the DORM database (stored in plain text) and read the output in R. This intermediate file containing the search results is deleted after being read in R (to plot and display the results on the user’s browser) to uphold the user’s privacy.

With large datasets, parallel computation, is known to improve performance (Nagurney 1996). Indeed, even with our limited four CPU-core (x86-64 architecture) setup, we observed performance improvements by using a parallelized version of the code. The gathered data indicated that parallelizing appropriate repetitive tasks with constructs such as `foreach::foreach()` and `parallel::mclapply()`, offer significant reductions in processing time (Supplementary Figure S2 F). The mclapply() implementations were the fastest methods across our range of tested number of operations (range: 600 – 60000) (Supplementary Figure S2 F). It is noteworthy that, in case of the foreach package, the gains in performance were dependent on the size of data, i.e., with smaller loops (600 operations) set significant overheads were incurred while setting up an environment for parallel processing (Supplementary Figure S2 F). Additionally, similar improvements were observed by parallizing the operations of saving and loading the standard .RDS file format in comparison to `base::saveRDS()` and `base::readRDS()` methods (Supplementary Figure S2 G-H).

### Contemporary databases are slow and resource intensive

One of the primary goals of DORM was to display the desired statistics and results faster than the contemporary databases like COSMIC, the ICGC data portal, cBioPortal and AACR GENIE data portal. To this end, Google Lighthouse was used to benchmark the performance of these databases and compare it to that of DORM to understand an end-user’s experience. DORM scored better than all the four databases (mean score for DORM: 84.7, COSMIC: 43.2, ICGC: 24.5, cBioPortal: 21, GENIE: 20.83) (Figure 1 A). In addition to taking the lowest time to become fully interactive (Figure 1 B), DORM was the only database that had zero seconds of blocking time (i.e., DORM remained responsive to user input) (Figure 1 C). Regarding client-side memory management, DORM had the lowest peak RAM usage (Figure 1 D), and the lowest RAM usage after garbage collection (i.e., after the browser was left to idle for 2 minutes) (Supplementary Figure S3 A) among the tested databases. Lighthouse performance score is a weighted mean of six individual parameters, namely, First Contentful Paint, Speed Index, Largest Contentful Paint, Time to interactive, Total Blocking Time, Cumulative Layout Shift, (details described in the materials & methods section titled “Benchmarking and testing performance”). The individual observations are plotted in Figure 1 B-C and Supplementary Figure S3 B-E.

**Figure 1.**
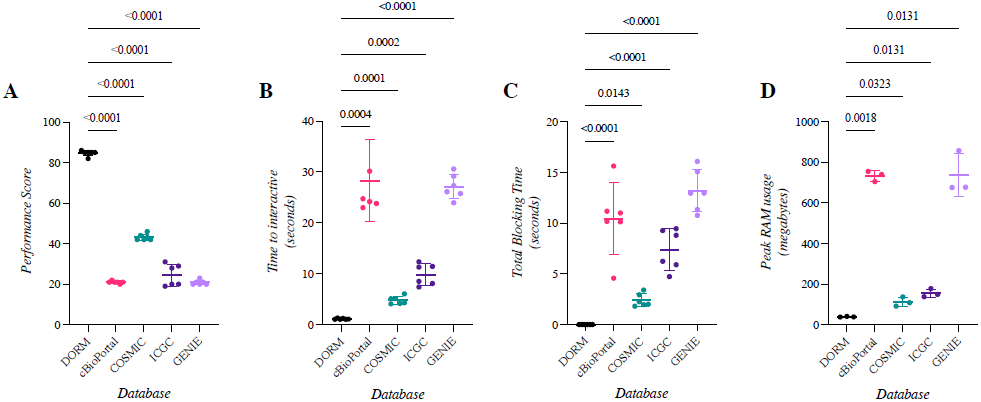
Comparison of databases and the performance of their websites. A search for EGFR mutations was done on each database, and the individual links for that search were used to test the performance of the databases with Google Lighthouse running on Google Chrome web browser. Scatter plots indicating mean and standard deviation of 3 to 6 observations for A) Lighthouse performance score B) Time to interactive (*y*-axis in seconds) C) Total blocking time (*y*-axis in seconds) D) Peak memory (RAM) usage by the web pages (*y*-axis in Megabytes).

### Identifying recurrent coding mutations after eradicating duplicate entries

All the major cancer databases (like COSMIC, cBioPortal, GENIE) which source information from multiple institutions share the common problem of mutations being reported multiple times due to same samples being included in different publications and/or studies. To counter this rampant issue of duplicate entries, we devised a mutation ID by using a combination of sample name, the protein name and the amino acid change and used it to remove the entries with duplicate mutation IDs. From the list of genuine coding mutations, COSMIC v95 had 78,569 duplicates, constituting 1.60 % of the filtered coding mutations (Supplementary Figure S1). After filtering out the non-recurrent mutations (i.e., mutations with a tissue-agnostic population frequency of 1), the data consists of 1,887,757 recurrent mutations which are comprised of 673,033 individual mutations (Supplementary Figure S1).

Interestingly, some samples (*n* = 1,207) in the dataset exclusively harbor non-recurrent coding mutations (Supplementary Figure S4 A), and overall, these individual samples belong to cancers of hematopoietic and lymphoid tissues, kidney, and autonomic ganglia (Supplementary Figure S4 A) and have very few coding mutations (range: 1 to 43 mutations per sample) (Supplementary Figure S4 B). On average, out of the total number of mutations across the 39 tissues included in the analysis, 39.1% of the mutations were recurrent (i.e., tissue-agnostic population frequency >1) in nature (Supplementary Figure S4 C). The highest percentage of recurrent mutations could be found in penile cancers (92.6%, *n* = 1,195 mutations; sample size = 10), thyroid cancers (89.6%; *n* = 139,883 mutations; sample size = 989) and meningeal malignancies (56.1%; n = 2,401 mutations; sample size = 163) (Supplementary Figure S4 C). The largest number of non-recurrent mutations were in cancers of the skin, large intestine, and lungs and together these contribute 48.5% of all the non-recurrent mutations. However, this was expected, as the samples from these three tissues also contribute a total of 46.6% of all the coding mutations in the dataset (Supplementary Figure S4 C).

The median mutational load (defined as the number of mutations per sample) in the dataset was 42 mutations/sample (mean (μ): 141, IQR: 81), with several outliers in different cancer types (Supplementary Figure S4 D). On average, samples from cancers of the endometrium (μ = 564), skin (μ = 508), and placenta (μ = 393) had the highest mutation load while samples from the autonomic ganglia (μ = 16), eye (μ = 16.8) and the adrenal gland (μ = 19) had the lowest (Supplementary Figure S4 D).

### Top recurrent mutations

Among the 100 most-recurrent mutations, the highest number of mutations are reported in TP53 (number of variants [*n*] = 19, frequency in cohort [ν] = 3,779), followed by KRAS (*n* = 9, ν = 3,006), PIK3CA (*n* = 6, ν = 1,808), BRAF (*n*=1, ν = 1,432), and NRAS (*n*=5, ν = 773) (Figure 2). Among these 100 recurrent mutations, 19 mutations (ν = 9,438) were in oncogenes, and 11 mutations (ν = 5,349) in tumor suppressor genes. The top three recurrent mutations were the amino acid substitutions BRAF V600E (ν = 1,432), KRAS G12D (ν = 977) and KRAS G12V (ν = 784) (Figure 2).

**Figure 2.**
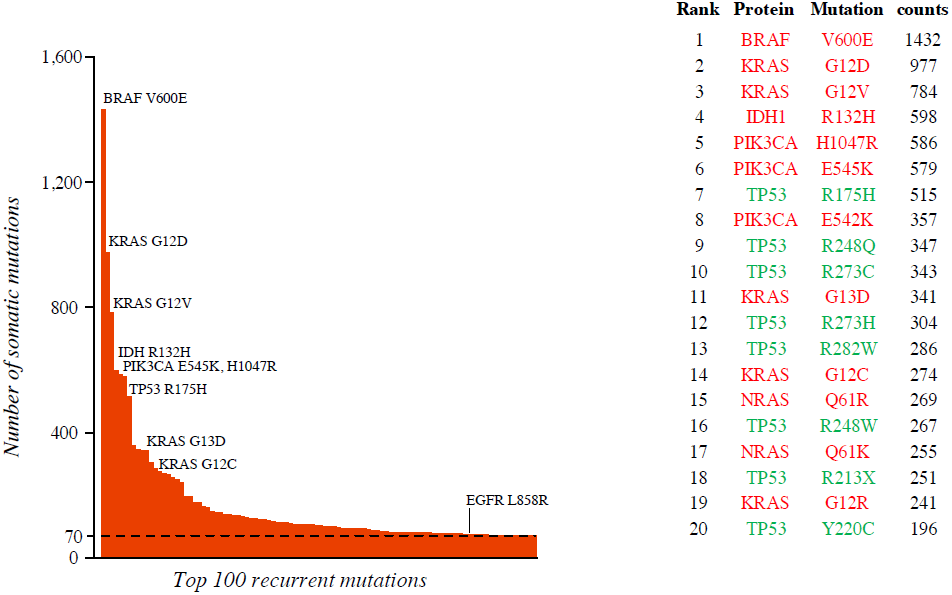
Distribution of top 100 recurrent mutations. Bar plots showing the top 100 most-frequently mutated proteins in the genome-wide somatic mutation data from COSMIC release v95. The top 20 mutations are listed in the table on the right, and the mutations in oncogenes are colored in red and the mutations in tumor suppressors are colored in green.

The survey of the cancer genomes, conducted here, was free from the selection bias that is introduced with targeted panels and selected sequencing. Although, while those are cost-effective strategies to identify driver and predictive mutations for cancers of selected histologies, they cause an over-representation of genes (and mutations in those genes) that are included in the selected panels which masks the true frequency of mutations in tumor tissues. This disparity was evident by the fact that EGFR L858R, the hotspot driver mutation in lung adenocarcinoma, ranked 84^th^ (ν = 75) in the list of most frequent amino acid substitutions (Figure 2). By contrast, it ranks 6th (ν = 10,631) when data from the targeted screens are also incorporated in computation of population frequencies.

### Website to browse the recurrent mutations

The processed database is hosted on a web server at the University of Turku and can be accessed at the URL https://eleniuslabtools.utu.fi/tools/DORM/Mutations (Figure 3). On receiving a connection request, the Shiny (https://shiny.rstudio.com/) web server spawns a new instance and displays the 50 mutations with the highest rate of recurrence. At the top of a page, there is a plot panel that consists of two dynamic plots that are updated in real-time in response to the user’s search queries. The bar plot on the left shows the cumulative frequency of the individual recurrent mutations in the population (Figure 3 A). The bar plot on the right shows the 25-most frequently mutated proteins across all the samples for the selected tissue (Figure 3 B). The plot is rendered as a high-resolution image in the user’s web browser in accordance with the browser’s dimensions and can be saved as an image straight from the browser. Query term(s) can be entered in the search bar (Figure 3 C), which updates the results in the table (Figure 3 D) showing the protein, the mutation, the aggregate frequency in the population, and the frequencies categorized by the primary site of the cancer. The results displayed in the table can be readily copied to a spreadsheet. Next to the search bar, there is a dropdown menu (Figure 3 E) to limit the number of results displayed in the table and the plot. The search or browsing can be restricted to a particular tissue from the menu (Figure 3 F). In addition to a button to reset the website and various parameters to their default value (Figure 3 G), there is a button to generate a direct link to a particular search (Figure 3 H). Clicking this button opens a dialog box (shown in Figure 3 I), with the link that can be used to perform the same search with the exact selected parameters. Clicking this button saves the search term(s) and the set parameter(s) anonymously on our server (i.e., no identifiable information is stored). Links like these facilitate sharing of the results, in addition to making it easy to repeat a search without having to enter the terms and set the parameters manually.

**Figure 3.**
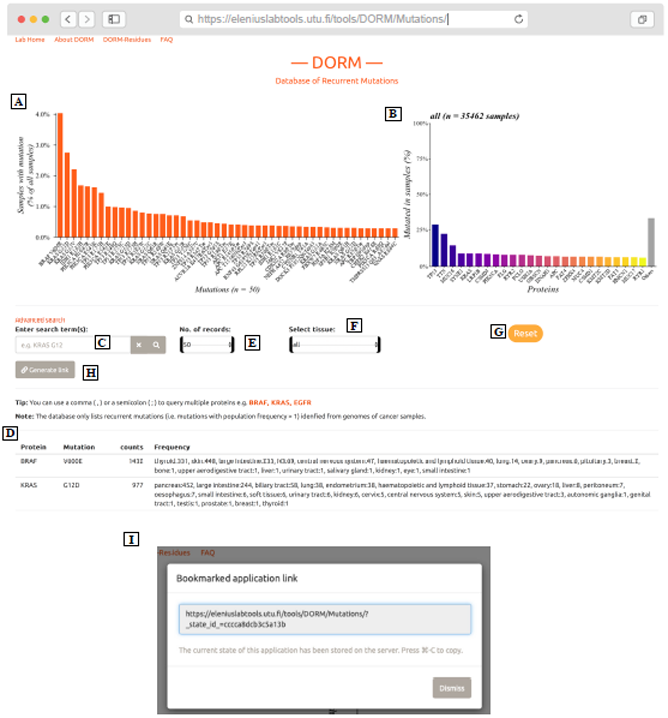
User interface for DORM : Database Of Recurrent Mutations. The default GUI of DORM, which hosted at https://eleniuslabtools.utu.fi/tools/DORM/Mutations/, shows the information about top 50 most-recurrent mutations identified from genomes of cancer samples. A) Dynamically updated bar plot, responds to search queries, and settings of dropdown menus in “E” and “F”. B) A bar chart showing the 25-most frequently mutated genes (color gradient) across all the samples in the selected tissue (can be changed from menu “F”). The “Others” bar represents the percentage of samples not containing mutations in any of the top 25 genes (bars with color gradient). C) The search bar can be used to query the database with several terms, as well as, regular expressions; an example is displayed in gray text. D) Table showing the protein name, mutation (displayed as amino acid change), the number of samples with that exact mutation, and the breakdown of the sample count by primary site of the cancer. E) Dropdown menu can change the number of records displayed in the table “D” and plotted in the bar plot “A”. F) Dropdown menu to limit the search to a specific tissue type. G) Button to reset the website and various parameters to their default values. H) Button to generate a direct link to repeat a search with the exact search terms and parameters. Clicking this opens the dialog box “I” which shows the link. I) Dialog box showing the direct link which can be used to conduct the exact search again without having to manually enter search term(s) and set the parameters.

### Searching the database

The user can search the database with terms such as “KRAS” (protein symbol), “V600E” (exact mutation), or “L858” (amino acid residue). The functions implemented in R and Shiny for searching the data were slow and unable to process a multi-term search query like “KRAS G12”. To facilitate this, a custom search function was written where a multi-term search query gets decomposed into constituent terms and the database is searched with GNU-grep (https://www.gnu.org/software/grep/) in a hierarchical manner. For instance, the server processes the above-mentioned search query by first shortlisting all the mutations in KRAS, then selecting only the mutations at the residue Gly 12. The search results are subsequently read and processed in R. Simultaneously, the plot panels are redrawn to correspond to these results and the user’s browser is updated with the search results. Depending on the search complexity and the number of results to be displayed, all of this computation happens within fractions of a second and the results are transmitted securely (TLS encryption) to the user’s web browser.

Circumstantially, when the search results consist of just a single protein (like “KRAS G12” mentioned above), the plot in the right panel changes to a bar plot showing the distribution of the tissues harboring the specific mutation(s) (Figure 4 A) that are displayed in the table. On the other hand, when a search result contains several proteins, the plot in the right panel changes to a pie showing the distribution of mutations in those proteins (Figure 4 B).

**Figure 4.**
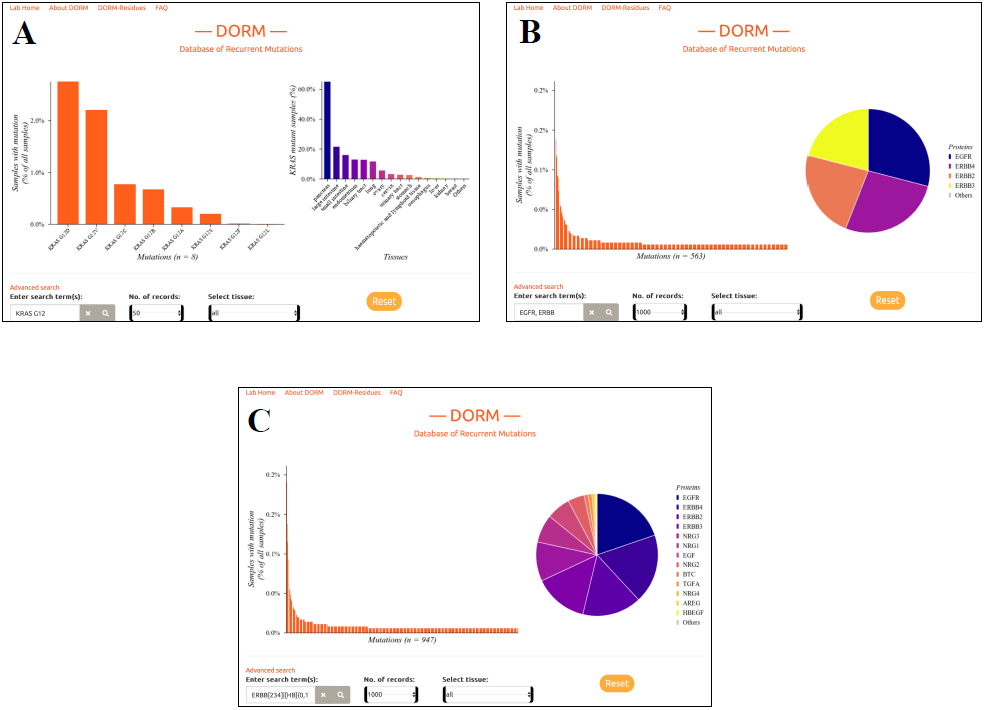
Dynamic plots in response to search results. Based on the context of the search results, DORM automatically generates these specific plots. A) If the search results (bar plot on the left) consists of just one protein (example results shown here for the query term `KRAS G12`), the bar plot on the right is updated to show the frequency of mutations in that protein in cancers of various tissues (*x*-axis). B) If the search results contain multiple proteins (query: `EGFR, ERBB`), a pie chart is displayed with the slices in the pie representing the proportion of mutations that can be attributed to the individual proteins. C) The distribution of search results when using a regular expression to search the database: `ERBB[234]|[HB]{0,1}EGF[R]{0,1}\>|NRG[1-4]|\<EP[GR]\>|AREG|BTC|TGFA.` This regular expression matches the four receptors and eleven ligands in the Epidermal Growth Factor Receptor family of proteins.

### Advanced search using regular expressions

The search bar (shown in Figure 3 C) in the DORM database supports advanced search using regular expressions. Regular expressions are an important tool in computational and data science and have been around since their inception in the 1950s (Kleene 1956). Regular expressions are a sequence of characters that define a set of strings and are a core component of almost all modern programming languages. In bioinformatics, regular expressions have been used in a myriad of diverse applications like establishing databases (Bairoch, Bucher and Hofmann 1997), determination of motifs from aligned protein sequences (Huang 2001), and in performing multiple sequence alignments (Arslan 2005). The language of regular expressions is known to have several dialects (i.e., syntax) (Zheng *et al*. 2021), and DORM supports the UNIX POSIX style regular expressions (IEEE and The Open Group 2018). A brief description and examples are presented here:

1. The pipe, i.e., | symbol, can be used for *either-or clause*, e.g., ‘`ERBB|EGFR`’ lists mutations in proteins EGFR, ERBB2, ERBB3 and ERBB4.
2. The square brackets, i.e., `[ABC]` structure, can be used to specify inclusions, e.g., ‘`NRG[1-4]`’ matches only the four of the Neuregulin ligands (NRG1, NRG2, NRG3, and NRG4). On the other hand, if we want to exclude results that can be done with `[^abc]` construct, e.g., ‘`ERBB[^4]`’ only matches ERBB2 and ERBB3 among the ERBB proteins leaving out ERBB4.
3. Word boundaries can be set with `\<` and `\>` operators, e.g., ‘`RAS\>`‘ matches all the proteins ending in ‘RAS’.
4. Match length modifying operators like `*`, `?`, +, and `{m,n}` are used to specify zero or more, at most one, at least one, or repetition for at least m times and at most n times respectively.

With the help of these regular expression operators one can formulate complex queries; for instance, if we are interested in searching the entire EGFR family of proteins (Yarden 2001) with one query, we can use a specific query like this:

‘`ERBB[234]|[HB]{0,1}EGF[R]{0,1}\>|NRG[1-4]|\<EP[GR]\>|AREG|BTC|TGFA`’

In the human protein repertoire, this regular expression matches the four receptors from the ERBB-family EGFR, ERBB2, ERBB3, ERBB4, and their eleven ligands AREG, BTC, TGFA, NRG1, NRG2, NRG3, NRG4, EGF, HBEGF, EPG and EGR; and nothing more (Figure 4 C).

## DISCUSSION

Next-generation sequencing (NGS) of cancer sample series has enabled more accurate understanding of cancer biology and helped to identify new predictive and therapeutic biomarkers. Here, we present DORM, a fast (Figure 1 and Supplementary Figure S3) and feature-rich (Table 1) web tool, which allows browsing its database that is derived from an unbiased analysis of somatic mutations identified by whole genome or whole exome NGS (consists of 91% of the coding mutations present in the COSMIC v95 data release). This strategy avoids encounters with the ill-effects of the selection biases that are introduced with the use of targeted NGS panels.

**Table 1.** Comparison of features between DORM and other public databases presenting somatic mutations identified from cancer samples. An asterisk indicates a caveat; which is clarified in the corresponding comment column.

| Searching & querying | DORM | COSMIC | ICGC | cBioportal | AACR Genie | Comment |
| --- | --- | --- | --- | --- | --- | --- |
| Protein | Yes | Yes | Yes | Yes | Yes |  |
| Individual Mutations (e.g. KRAS G12C) | Yes | Yes | Yes * | Yes * | Yes * | cBioPortal & Genie require searching for the gene and then the mutation. ICGC requires clicking the mutation from a list of results in a dropdown menu. |
| Protein sets | Yes | No | No | Yes | Yes |  |
| Tissues | Yes | Yes | Yes | No | No |  |
| Regular expression | Yes | No | No | No | No |  |
| Substring search (searching for RAS shows HRAS, KRAS, NRAS, etc.) | Yes | Yes | Yes | No | No |  |
| Additional features |  |  |  |  |  | Comment |
| Direct link to save and share search results | Yes | Yes | Yes | Yes | Yes |  |
| Summarize by Residue | Yes | No | No | No | No | cBioPortal Lollipop occasionally groups hotspot mutations at a residue as a single lollipop. |
| Duplicate samples | No | Yes | No | Yes | Yes |  |
| Display most recurrent mutations for a tissue or protein set | Yes | No | Yes | No | No |  |
| Show frequency of a protein being mutated in various tissues | Yes | Yes * | No | Yes | No | Possible on COSMIC Cancer Browser. |
| Number of rows displayed in table | 10-10000 | 10-100 | 10-50 | 25 | 25 |  |
| Free from cookies, trackers and/or analytics | Yes | No | No | No | No |  |

In addition to be performant and fast (Figure 1 and Supplementary Figure S3), DORM has several advantages (Table 1), most notably the ability to directly search for sets of proteins and use regular expressions. Additionally, DORM is the only database that can summarize the mutations at an amino acid residue level, (this feature is accessible by clicking “DORM-Residues” link at the top of the page) (direct link: https://eleniuslabtools.utu.fi/tools/DORM/Residues/). Data from all other databases requires manual processing to retrieve this information. DORM is also the only database to allow the user to view a large amount of the data without having to click through numerous pages of results (on DORM, users can choose from a range of 10-10000 results to display). DORM is also the only database that is free of cookies, trackers, and any embedded analytics.

COSMIC, cBioPortal and AACR Genie feature duplicate entries, while DORM and ICGC do not. Individual mutations, like KRAS G12C, can be directly searched on DORM as well as COSMIC. The cBioPortal, and the AACR GENIE (based on the cBioPortal user interface) do not offer the users to limit the search to specific tissues. On ICGC the implementation is similar to DORM (i.e., requires selection from a menu), but on COSMIC the user has to click the tissue from a table in the “Tissue distribution” section.

In the quest for speed and performance, certain compromises had to be made that constitute the limitations of DORM (Table 2). For instance, DORM does not incorporate or display the information about copy number variations or structural variations and has stripped all the detailed sample- and study-level information. Like some other databases in our comparison, DORM also doesn’t display fusions or non-coding mutations, allow selecting multiple tissues, and display a lollipop diagram which are a nice tool that place the mutations in context of the primary sequence of the proteins.

**Table 2.**
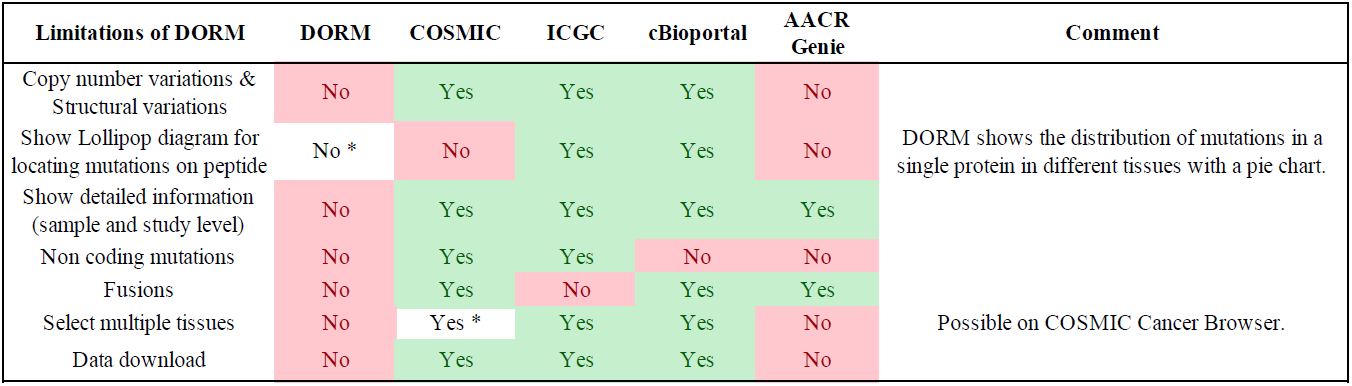
Limitations of DORM in comparison to other public databases presenting somatic mutations identified from cancer samples. An asterisk indicates a caveat; which is clarified in the corresponding comment column.

DORM is lightweight, and, by using our open-source codebase (see methods for links to the repositories), it can be run on normal consumer hardware. DORM is publicly available on a virtual private server that allows us to scale up the resources with an increase in demand. We believe that DORM can improve accessibility of the important information about recurrent mutations by being faster and by consuming lesser resources than the competition.

---

**Figure S1.**
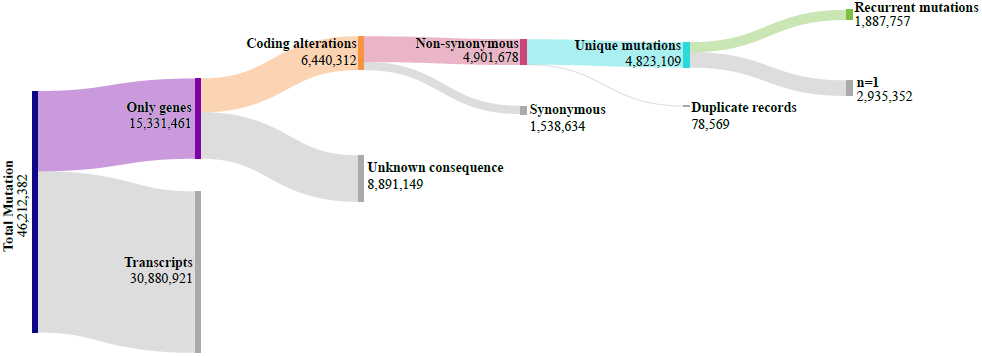
Filtering scheme to isolate recurrent mutations from COSMIC data. Out of 46 million mutations, 30.8 million mutations were removed as they can be attributed to duplicate transcripts of genes. 8.89 million mutations with unreported / unknown consequence were removed. 1.53 million silent mutations were removed, which do not produce any change in the protein. Subsequently, 78,569 duplicate records were removed as they are present due to incorporation of some samples in multiple studies. After this filtering process, 4.82 million unique coding mutations that remained were processed to create the DORM database which summarizes the 1.88 million recurrent mutations (mutations with tissue-agnostic population frequency >1).

**Figure S2.**
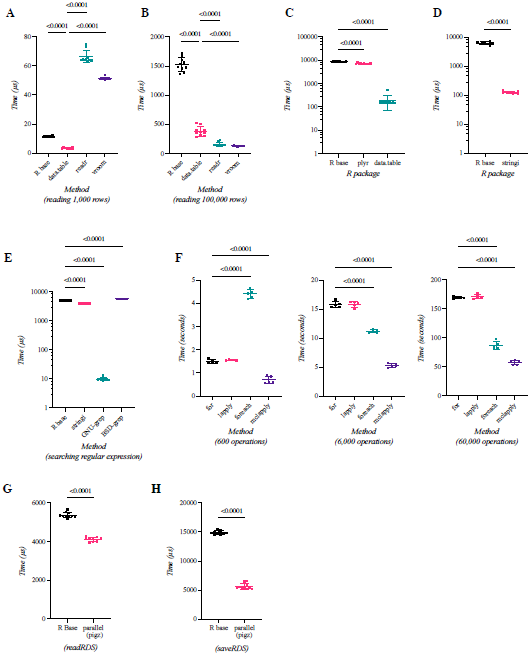
Comparison of strategies and methods for performing various computational operations. Scatter plots showing mean and standard deviation for A-B) Different R packages for reading a tabseparated value (TSV) file or 1000 rows (shown in A) and 100,000 rows (shown in B). C) Various approaches of generating a frequency table. D) Searching for a regular expression (pattern: `[ACDEFGHIKLMNPQRSTVWYX]?[0-9]+`) using R base and stringi package. E) Searching the data with a regular expression (pattern: `EGFR|ERBB[2-4]|[HKN]RAS\>`) using the indicated methods. For giving the functions in R the best chance, the data was preloaded in the R outside the code for timing the benchmark. F) Comparing various looping constructs for 600, 6000 and 60000 operations, for and lapply are serialized loops, foreach and mclapply are their parallel alternatives. G) Writing an R object containing a table (1 million rows) as an RDS file with either the R base serialized version or our custom parallelized version using ‘pigz’ for compression. H) Reading an RDS file (written with our parallel version of saveRDS) with the R base serialized version or our custom parallelized version using ‘pigz’ for decompression.

**Figure S3.**
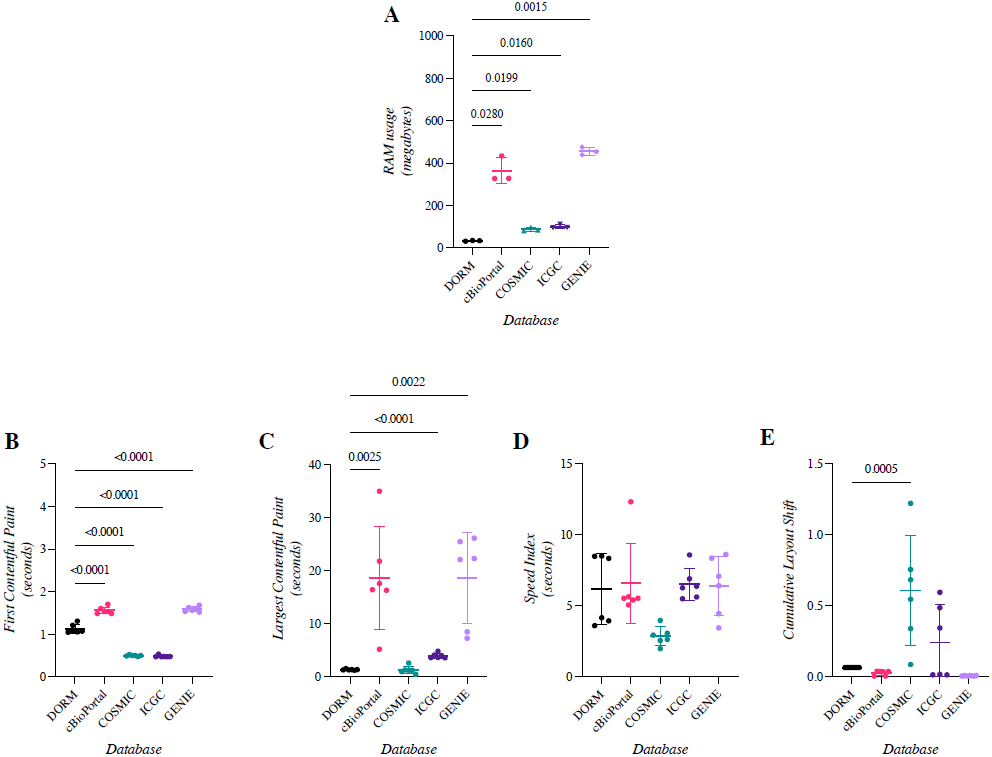
Comparison of databases and the performance of their websites. A search for EGFR mutations was done on each database, and the individual links for that search were used to test the performance of the databases with Google Lighthouse running on Google Chrome web browser. Scatter plots with mean and standard deviation for 3 to six observations for A) Memory (RAM) usage after garbage collection (browser idling for 2 minutes) while browsing the indicated databases (*y*-axis in Megabytes) B) First contentful paint (*y*-axis in seconds) C) Largest contentful paint (*y*-axis in seconds) D) Speed index (*y*-axis in seconds) E) Cumulative layout shift.

**Figure S4.**
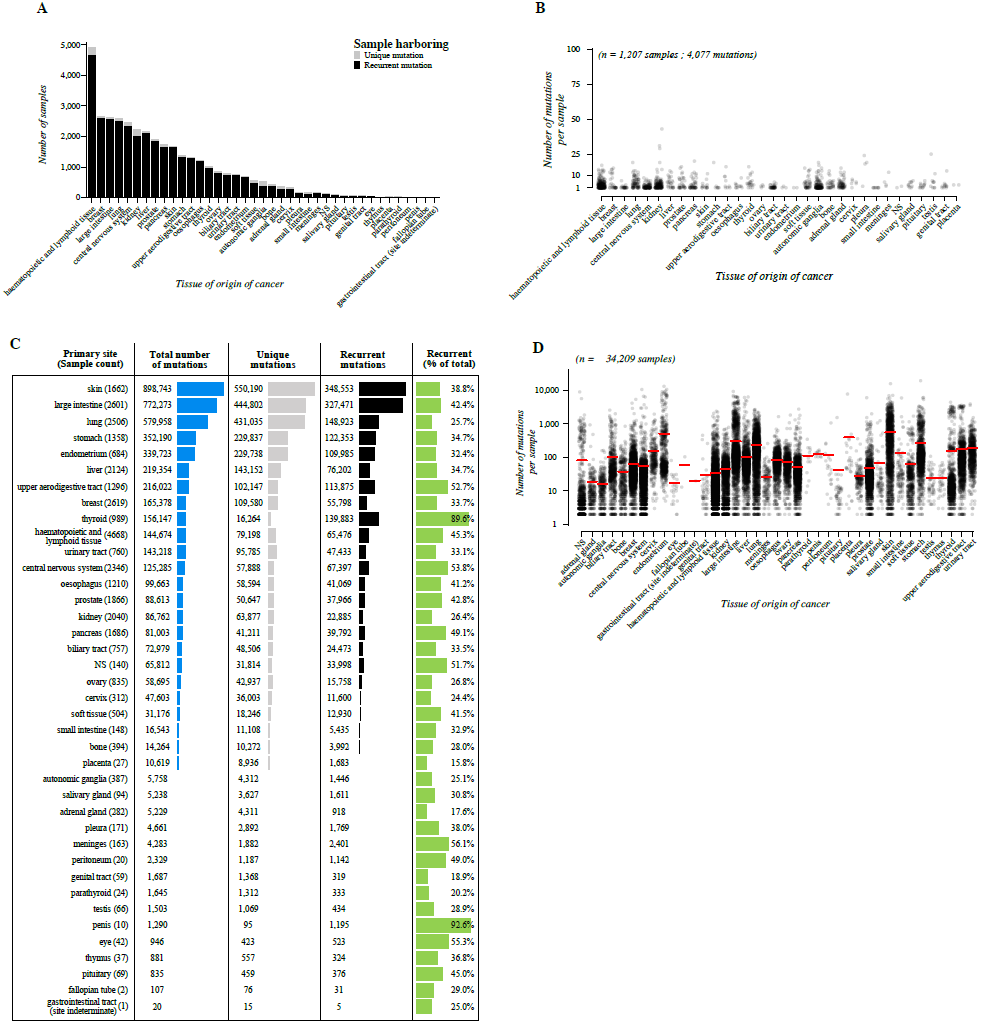
Distribution of mutations across sam les included in t e analysis. A) Bar chart showing the number of samples harboring at least one recurrent coding mutation, or samples having only unique coding mutations. Recurrent coding mutations are defined as protein sequence-altering mutations with a tissue-agnostic population frequency > 1. Samples are categorized by primary site of the cancer (*x*-axis). B) Dot plot showing the mutational load of individual samples (*n* = 1,207) harboring only unique coding mutations (*n* = 4,077) (i.e., samples comprising the gray part of the bars in panel A). Samples are categorized by primary site of the cancer (*x*-axis). C) Distribution of the individual coding mutations categorized by the primary site of the cancer, showing the total number of mutations, the number of unique (mutations never observed in any other sample (tissueagnostic) in the dataset) and recurrent mutations (tissue-agnostic population frequency > 1), and the proportion of the recurrent mutations from the total number of mutations are shown as a percentage. D) Dot plot showing distribution of mutational load (*y*-axis in log scale) in samples (*n* = 35,462). Each point represents a sample. Red horizontal line shows the mean mutational load for samples of different primary sites of the cancer (*x*-axis). NS = tissue is not specified.

## REFERENCES

Aho AV, Kernighan BW, Weinberger PJ. The AWK Programming Language. Addison-Wesley Publishing Company, 1988.

Arslan AN. Multiple Sequence Alignment Containing a Sequence of Regular Expressions. 2005 IEEE Symposium on Computational Intelligence in Bioinformatics and Computational Biology. La Jolla, CA, USA: IEEE, 2005, 1–7.

Bairoch A, Bucher P, Hofmann K. The PROSITE database, its status in 1997. Nucleic Acids Res 1997;25:217–21.

Benjamini Y, Krieger AM, Yekutieli D. Adaptive linear step-up procedures that control the false discovery rate. Biometrika 2006;93:491–507.

Campbell PJ, Getz G, Korbel JO et al. Pan-cancer analysis of whole genomes. Nature 2020;578:82–93.

Cerami E, Gao J, Dogrusoz U et al. The cBio Cancer Genomics Portal: An Open Platform for Exploring Multidimensional Cancer Genomics Data. Cancer Discov 2012;2:401–4.

Chang MT, Asthana S, Gao SP et al. Identifying recurrent mutations in cancer reveals widespread lineage diversity and mutational specificity. Nat Biotechnol 2016;34:1–11.

Chang W, Cheng J, Allaire JJ et al. Shiny: Web Application Framework for R., 2021. Dowle M, Srinivasan A. Data.Table: Extension of ‘data.Frame’., 2021.

Gagolewski M. stringi: Fast and portable character string processing in R. J Stat Softw 2021.

Gao J, Aksoy BA, Dogrusoz U et al. Integrative analysis of complex cancer genomics and clinical profiles using the cBioPortal. Sci Signal 2013;6:pl1.

Hauschild A, Grob J-J, Demidov LV et al. Dabrafenib in BRAF-mutated metastatic melanoma: a multicentre, open-label, phase 3 randomised controlled trial. The Lancet 2012;380:358–65.

Hong DS, Fakih MG, Strickler JH et al. KRAS G12C Inhibition with Sotorasib in Advanced Solid Tumors. N Engl J Med 2020;383:1207–17.

Huang JY. The EMOTIF database. Nucleic Acids Res 2001;29:202–4. IEEE, The Open Group. The Open Group Base Specifications Issue 7. 2018.

Kleene SC. Representation of events in nerve nets and finite automata. Autom Stud 1956;34:3–41.

Lynch TJ, Bell DW, Sordella R et al. Activating Mutations in the Epidermal Growth Factor Receptor Underlying Responsiveness of Non–Small-Cell Lung Cancer to Gefitinib. N Engl J Med 2004;350:2129–39.

Martincorena I, Campbell PJ. Somatic mutation in cancer and normal cells. Science 2015;349:1483–9. Mersmann O. Microbenchmark: Accurate Timing Functions., 2021.

Nagurney A. Chapter 7 Parallel computation. Vol 1. Elsevier, 1996, 335–404.

National Institute of Standards and Technology. Advanced Encryption Standard (AES). Gaithersburg, MD: National Institute of Standards and Technology, 2001:NIST FIPS 197.

Ooms J. The jsonlite Package: A Practical and Consistent Mapping Between JSON Data and R Objects. ArXiv14032805 StatCO 2014.

Paez JG, Jänne P, Lee JC et al. EGFR mutations in lung cancer: correlation with clinical response to gefitinib therapy. Science 2004;304:1497–500.

R Core Team. R: A Language and Environment for Statistical Computing. Vienna, Austria: R Foundation for Statistical Computing, 2018.

Rescorla E. The Transport Layer Security (TLS) Protocol Version 1.3. RFC Editor, 2018.

Tate JG, Bamford S, Jubb HC et al. COSMIC: the Catalogue Of Somatic Mutations In Cancer. Nucleic Acids Res 2019;47:D941–7.

The AACR Project GENIE Consortium. AACR Project GENIE: Powering Precision Medicine through an International Consortium. Cancer Discov 2017;7:818–31.

Watson JD, Baker TA, Bell SP et al. Molecular Biology of the Gene. 6th ed. San Francisco; Cold Spring Harbor, N.Y.: Pearson/Benjamin Cummings; Cold Spring Harbor Laboratory Press, 2007.

Yarden Y. The EGFR family and its ligands in human cancer: signalling mechanisms and therapeutic opportunities. Eur J Cancer Oxf Engl 1990 2001;37 Suppl 4:S3-8.

Zhang J, Baran J, Cros A et al. International Cancer Genome Consortium Data Portal--a one-stop shop for cancer genomics data. Database J Biol Databases Curation 2011;2011:bar026.

Zheng L-X, Ma S, Chen Z-X et al. Ensuring the Correctness of Regular Expressions: A Review. Int J Autom Comput 2021;18:521–35.

